# Circadian Clock Core Component Bmal1 Dictates Cell Cycle Rhythm of Proliferating Hepatocytes during Liver Regeneration

**DOI:** 10.1101/2021.08.11.455958

**Authors:** Huaizhou Jiang, Veronica Garcia, Jennifer Abla Yanum, Joonyong Lee, Guoli Dai

## Abstract

Following partial hepatectomy (PH), the majority of remnant hepatocytes synchronously enter and rhythmically progress through the cell cycle for three major rounds to regain lost liver mass. Whether and how the circadian clock core component Bmal1 modulates this process remains elusive. We performed PH on *Bmal1*^+/+^ and hepatocyte-specific *Bmal1* knockout (*Bmal1*^hep-/-^) mice and compared the initiation and progression of the hepatocyte cell cycle. After PH, Bmal1^+/+^ hepatocytes exhibited three major waves of nuclear DNA synthesis. In contrast, in *Bmal1*^hep-/-^hepatocytes, the first wave of nuclear DNA synthesis was delayed by 12 h, and the third such wave was lost. Following PH, Bmal1^+/+^ hepatocytes underwent three major waves of mitosis, whereas *Bmal1*^hep-/-^hepatocytes fully abolished mitotic oscillation. These Bmal1-dependent disruptions in the rhythmicity of hepatocyte cell cycle after PH were accompanied by suppressed expression peaks of a group of cell cycle components and regulators, and dysregulated activation patterns of mitogenic signaling molecules c-Met and EGFR. Moreover, *Bmal1*^+/+^ hepatocytes rhythmically accumulated fat as they expanded following PH, whereas this phenomenon was largely inhibited in *Bmal1*^hep-/-^hepatocytes. In addition, during late stages of liver regrowth, Bmal1 absence in hepatocytes caused the activation of redox sensor Nrf2, suggesting an oxidative stress state in regenerated liver tissue. Collectively, we demonstrated that during liver regeneration, Bmal1 partially modulates the oscillation of S-phase progression, fully controls the rhythmicity of M-phase advancement, and largely governs fluctuations in fat metabolism in replicating hepatocytes, and eventually determines the redox state of regenerated livers.

**New and Noteworthy:** We demonstrated that Bmal1 centrally controls the synchronicity and rhythmicity of the cell cycle and lipid accumulation in replicating hepatocytes during liver regeneration. Bmal1 plays these roles, at least in part, by ensuring formation of the expression peaks of cell cycle components and regulators, as well as the timing and levels of activation of mitogenic signaling molecules.

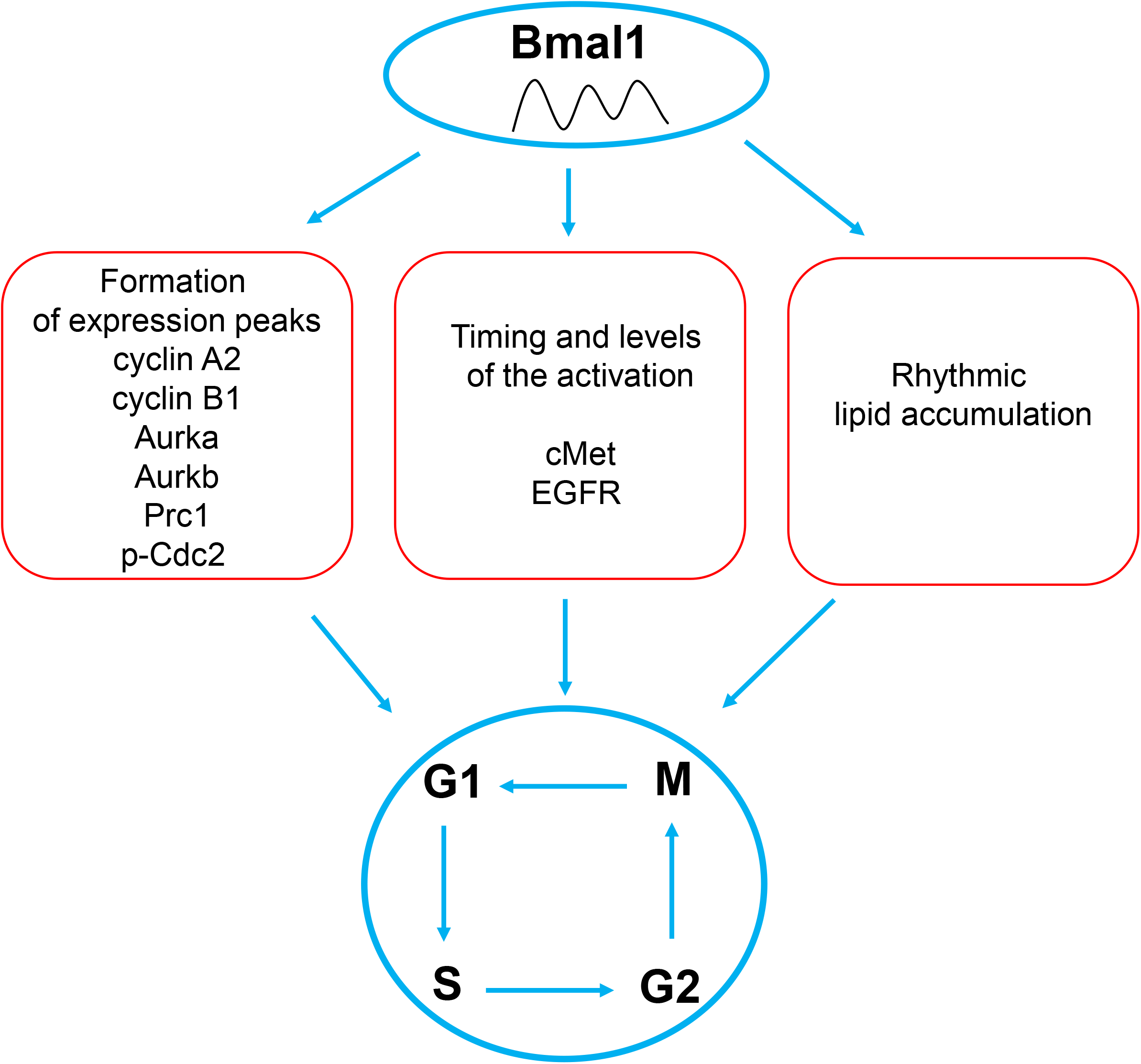
Graphical Abstract.

## Introduction

Following liver resection or acute liver injury, remnant hepatocytes synchronously enter the cell cycle and continuously undergo three major waves of replication to restore the original liver mass(42). This process is driven by the highly coordinated activation of a variety of extrahepatic and intrahepatic signals and pathways, which orchestrate the timely and dynamic expression of cell cycle regulators and components(22). However, how the synchronicity and rhythmicity of cell cycle progression is regulated remains elusive.

The core component of the mammalian circadian clock is composed of Bmal1 and Clock. These two transcription factors form a heterodimer, operate a set of transcriptional/translational feedback loops, generate diurnal outputs of the cellular transcriptome, and ultimately govern various rhythmic biological processes (3, 7). Of note, the circadian clock interacts with the cell cycle to regulate tissue renewal and repair(30). Several lines of evidence, including our previous work, support the notion that the circadian clock controls the timing of hepatocyte mitosis during liver regeneration(20, 33, 42). However, it remains unclear whether and how Bmal1 plays such a role. We aimed to determine how loss of Bmal1 function affects the initiation and progression of the cell cycle in replicating hepatocytes during the entire period of liver regrowth after partial hepatectomy (PH).

## Materials and Methods

### Animal care and use

Hepatocyte-specific *Bmal1* knockout mice were generated by crossing Bmal1^flox/flox^ mice (stock# 009100, The Jackson Laboratory, Bar Harbor, ME, USA) with Albumin-Cre mice (stock# 003574, The Jackson Laboratory). Mice were housed in plastic cages at 22 ± 1°C on a 12-hour light/12-hour dark cycle with lights on from 6:00 am to 6:00 pm. Standard rodent chow and water were provided *ad libitum* throughout the feeding period. Male mice (3-4 months old) were subjected to standard two-thirds PH with intact gall bladders following a procedure described previously(6, 12). The surgery was performed in the morning between 10:00 am and 12:00 pm, with consistent surgical time on all mice(20). BrdU was injected (100 mg/kg, i.p.) 2 h before euthanization at each time point. The isolated livers were weighed. Liver pieces were fixed in 10% neutral buffered formalin, embedded in optimal cutting temperature compound (Tissue Tek, Torrance, CA, USA), or frozen in liquid nitrogen for various subsequent analyses. All animal experiments were conducted in accordance with the National Institutes of Health Guide for the Care and Use of Laboratory Animals. Protocols for the care and use of animals were approved by the Indiana University-Purdue University Indianapolis Animal Care and Use Committee.

### Histology and Immunohistochemistry

Formalin-fixed and paraffin-embedded liver sections were subjected to Ki-67, BrdU, or p-histone H3 immunostaining to visualize and count proliferating hepatocytes. Primary antibodies against Ki-67 (RM-9106, Thermo Fisher Scientific, Waltham, MA, USA), BrdU (#5292, Cell Signaling Technology, Danvers, MA, USA), or p-histone H3 (#9764, Cell Signaling Technology) were used for immunostaining. Ki67-, BrdU-, or p-histone H3-positive hepatocytes were counted in five randomly chosen microscopic fields per section at 200× magnification using Image-Pro Plus software (Media Cybernetics, Rockville, MD, USA). Oil Red O (Sigma-Aldrich Co., St. Louis, MO, USA) staining was performed on frozen liver sections in order to visualize intracellular lipid droplets. Stained areas were analyzed in five randomly chosen microscope fields at 200× magnification per section using Image-Pro Plus software.

### Western Blot Analysis

Liver homogenates (10 or 30 µg) were separated by polyacrylamide gel electrophoresis under reducing conditions. Proteins from the gels were electrophoretically transferred onto polyvinylidene difluoride (PVDF) membranes. Antibodies against cyclin D1 (#2922), cyclin B1 (#4138), and glyceraldehyde 3-phosphate dehydrogenase (GADPH) (#5174) were purchased from Cell Signaling Technology; Bmal1 (SC-373955), Wee1 (SC-9037), p-Cdc2 p34 (Tyr^15^) (SC-7898), and Cdc2 p34 (SC-54) from Santa Cruz Biotechnology, Dallas, TX, USA; cyclin A2 (ab32386), c-Met (ab47431), p-cMet (ab5662), and NQO1 (ab80588) from Abcam, Cambridge, MA, USA and EGFR (#1114-1) and p-EGFR (#1139) from Epitomics (Burlingame, CA, USA). Immune complexes were detected using an enhanced chemiluminescence system (Pierce, Rockford, IL, USA).

### Quantitative real-time polymerase chain reaction

Total RNA was isolated from frozen liver tissue using TRIzol reagent according to the manufacturer’s protocol (Invitrogen, Carlsbad, CA, USA). cDNA was synthesized from total RNA (1 µg) of each sample using a Verso cDNA Kit (Thermo Fisher Scientific), diluted 4-fold with water, and subjected to RT-qPCR to quantify mRNA levels. TaqMan Universal PCR Master Mix, PCR primers and TaqMan MGB probes specific for mouse *Bmal1* (Mm01269610_m1), *Dbp* (Mm00497539_m1), *cyclin A2* (Mm00438063_m1), *cyclin B1* (Mm03053893_gh), *Aurka* (Mm01248177_m1), *Aurkb*, (Mm01718146_gl), *Prc1* (Mm01320564_m1), *Wee1* (Mm00494175_m1), *Per1* (Mm00501813_m1), *Per2* (Mm00478099_m1), *Elovl6* (Mm00851223_s1), *Fads1* (Mm00507605_m1), *Fads2* (Mm00517221_m1), *Srebf1* (Mm00550338_m1), *Ppara* (Mm00440939_m1), *Pparg* (Mm00440940_m1), *Cd36* (Mm00432403_m1), *Fasn* (Mm00662319_m1), *Cry1* (Mm00514392_m1), *Cry2* (Mm00546062_m1), and *Ppia* (Mm02342430_gl) were purchased from Applied Biosystems (Foster City, CA, USA). RT-qPCR amplification reactions were carried out using the ABI Prism 7900 sequence detection system (Applied Biosystems) with an initial hold step (50°C for 2 min followed by 95°C for 10 min) and 40 cycles of a 2-step PCR thermocycling protocol (92°C for 15 s and 60°C for 1 min). The comparative C_T_ method was used for relative quantification of the amount of mRNA in each sample normalized to *Ppia* transcript levels.

### Statistical analysis

Data are shown as means ± standard deviation (SD). Statistical analyses were performed using one-way analysis of variance. Comparisons of means were determined using post-hoc analysis. Significant differences were defined as *p* < 0.05.

## Results

### Hepatocyte-specific *Bmal1* knockout disrupted S-phase progression and abolished mitotic waves in replicating hepatocytes in regenerating livers

For simplicity, we define *Bmal1*^+/+^;*Albumin-Cre*^+^ mice as *Bmal1*^+/+^ mice and *Bmal1*^flox/flox^ ;*Albumin-Cre*^+^ mice as hepatocyte-specific *Bmal1* knockout (*Bmal1*^hep-/-^) mice.

We performed PH on *Bmal1*^+/+^ and *Bmal1*^hep-/-^mice and subsequently conducted the following assessments at various time points within a 7-day period. We first quantified Ki67-positive hepatocytes in regenerated liver tissue. Because Ki67 is expressed in all phases of the cell cycle, we used this marker to estimate the total number of cycling hepatocytes. We observed Bmal1-dependent alterations in this parameter at multiple time points after PH (Fig. 1). These data indicated that Bmal1 deficiency in hepatocytes affects PH-induced hepatocyte proliferation.

**Figure 1.**
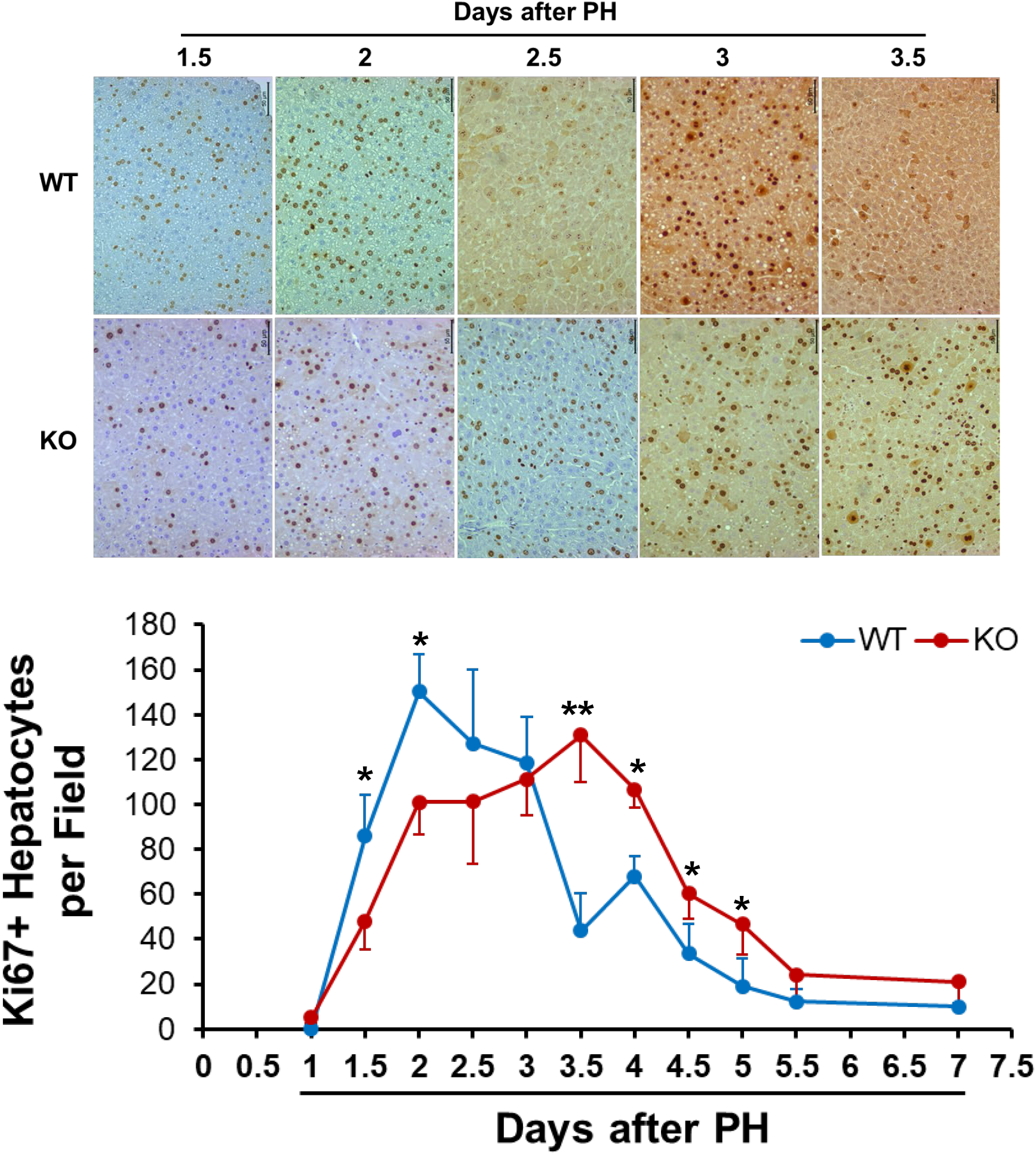
Numbers of cycling hepatocytes after partial hepatectomy (PH). *Bmal1*^+/+^ (WT) and hepatocyte-specific *Bmal1* knockout (KO) 3-4-month-old male mice were subjected to PH and sacrificed at the indicated time points. Ki-67 immunostaining was performed with liver sections. **(Top panel)** Representative liver sections showing Ki67-positive hepatocytes. **(Bottom panel)** Ki67-positive hepatocytes were counted at 200× magnification in five randomly chosen fields per section. The results are shown as mean numbers per field ± SD (n = 5 mice/time point/genotype; *, *p* < 0.05; **, *p* < 0.01).

To gain insight into the S-phase progression of proliferating hepatocytes, we injected bromodeoxyuridine (BrdU) into mice 1 h before each time point and quantified BrdU-positive (S-phase) hepatocytes (Fig. 2). As expected, *Bmal1*^+/+^ regenerating livers underwent three major waves of DNA synthesis, which consecutively occurred and progressively weakened at days 1.5, 3, and 4 post-PH. In contrast, in *Bmal1*^hep-/-^regenerating livers, the first wave of DNA synthesis was delayed by 12 h, the second wave occurred in a timely manner, and the third wave disappeared. Cycling *Bmal1*^hep-/-^hepatocytes exhibited dysregulated S-phase progression following PH.

**Figure 2.**
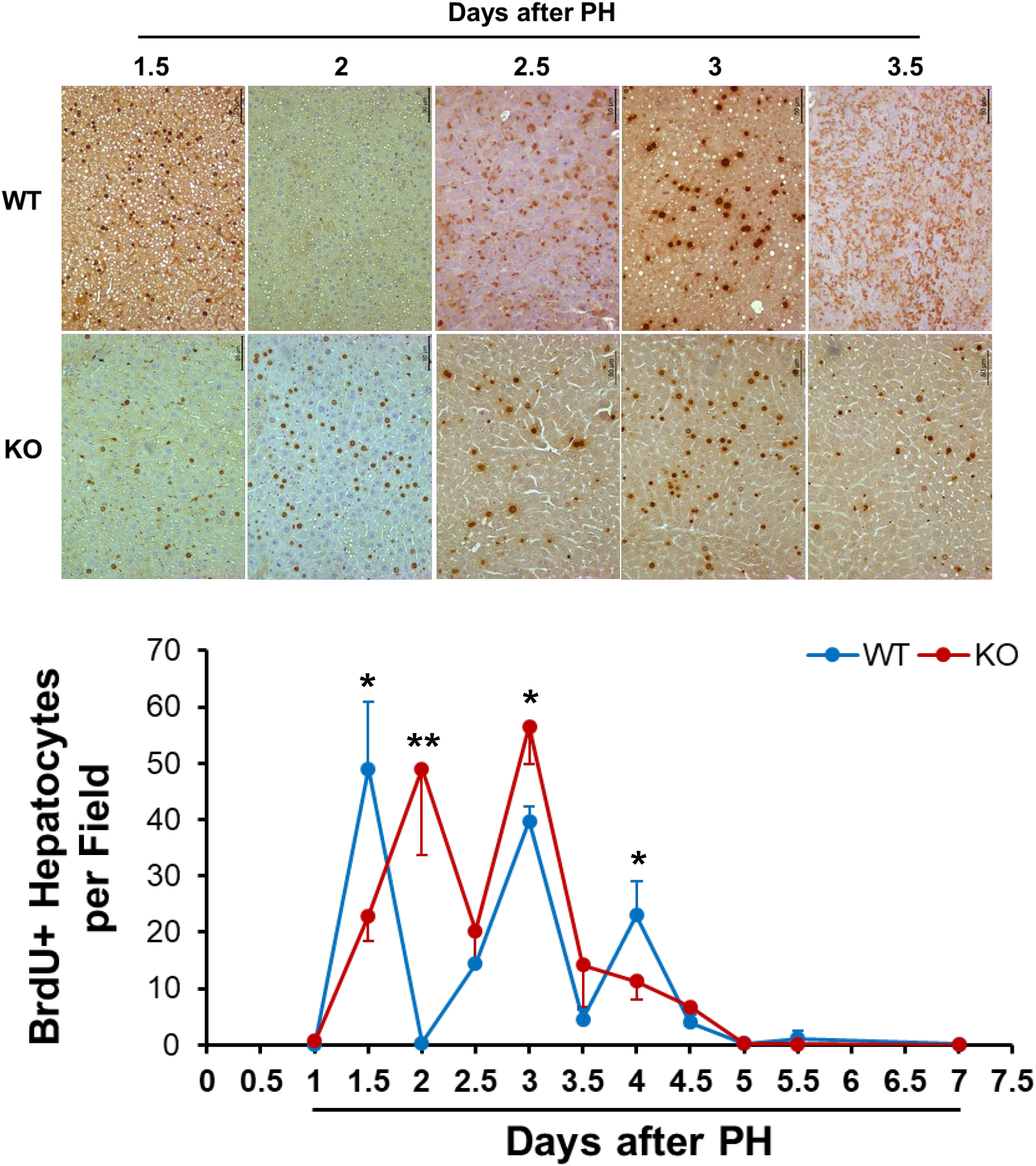
Numbers of hepatocytes undergoing DNA synthesis after PH. *Bmal1*^+/+^ (WT) and hepatocyte-specific *Bmal1* knockout (KO) 3-4-month-old male mice were subjected to PH and sacrificed at the indicated time points. One hour prior to sacrifice, BrdU was injected (100 mg/kg, i.p.). Liver sections were subjected to BrdU immunostaining. **(Top panel)** Representative liver sections showing BrdU-positive hepatocytes. **(Bottom panel)** BrdU-positive hepatocytes were counted at 200× magnification in 5 randomly chosen fields per section. The data are shown as the mean numbers per field ± SD (n = 5 mice/time point/genotype; *, *p* < 0.05; **, *p* < 0.01).

We next examined the M-phase progression of replicating hepatocytes by quantifying hepatocytes positive for p-histone H3, an M-phase cell cycle marker (Fig. 3). As anticipated, *Bmal1*^+/+^ regenerating hepatocytes displayed three major waves of mitosis, which were sequentially formed and gradually reduced at days 2, 3, and 4 after PH. In sharp contrast, *Bmal1*^hep-/-^regenerating hepatocytes completely lost the oscillation in M-phase progression. Even on day 1.5 following PH, when *Bmal1*^+/+^ cycling hepatocytes were undergoing the first S phase (Fig. 2), a significant number of cycling *Bmal1*^hep-/-^hepatocytes had already entered M phase. Thus, Bmal1 is essential for the formation of hepatocyte mitotic waves during PH-induced liver regrowth.

**Figure 3.**
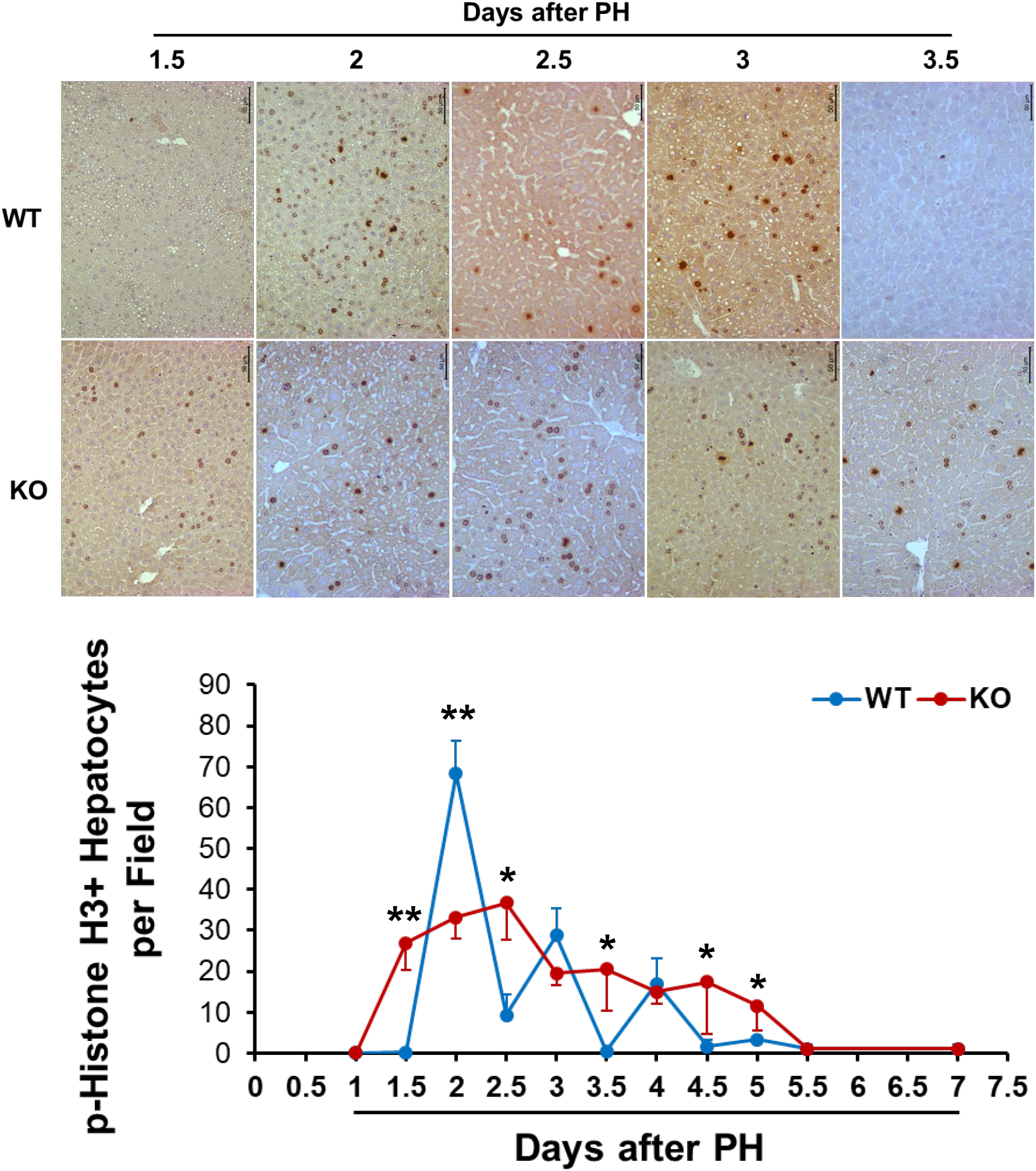
Numbers of hepatocytes undergoing mitosis after partial hepatectomy (PH). *Bmal1*^+/+^ (WT) and hepatocyte-specific *Bmal1* knockout (KO) 3-4-month-old male mice were subjected to PH and sacrificed at the indicated time points. Liver sections were immunostained with anti-p-Histone H3 antibody. **(Top panel)** Representative liver sections show p-Histone H3-positive hepatocytes. **(Bottom panel)** p-Histone H3-positive hepatocytes were counted at 200× magnification in five randomly chosen fields per section. The data are shown as means per field ± SD (n = 5 mice/time point/genotype; *, *p* < 0.05; **, *p* < 0.01).

### Hepatocyte-specific *Bmal1* knockout results in reduced fat accumulation in regenerating livers

It is known that, after PH, replicating hepatocytes dynamically accumulate fat, which is coupled to cell cycle progression (42). We conducted Oil Red O staining and measured the staining areas in the two genotypes of regenerating livers (Fig. 4). As a result, *Bmal1*^+/+^ regenerating livers showed massive fat deposits at days 1, 1.5, 3, and 4 following PH. In contrast, *Bmal1*^hep-/-^regenerating livers displayed reduced fat staining at day 1 post-PH, and largely diminished fat content thereafter. Therefore, regenerating hepatocytes require Bmal1 to dynamically govern lipid metabolism.

**Figure 4.**
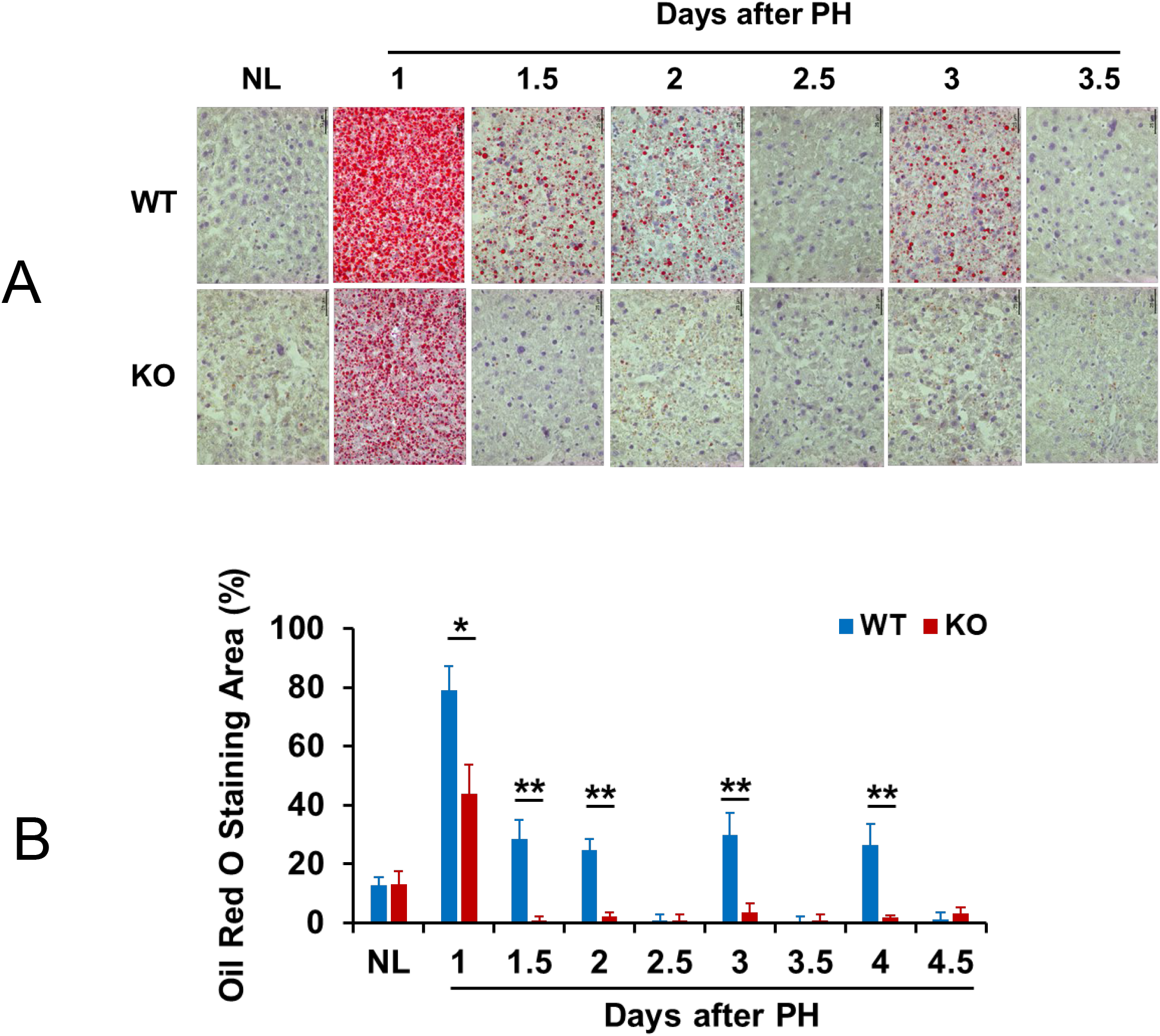
Fat accumulation in regenerating livers. *Bmal1*^+/+^ (WT) and hepatocyte-specific *Bmal1* knockout (KO) 3-4-month-old male mice were subjected to PH and sacrificed at the indicated time points. Frozen liver sections were prepared and used for Oil red O staining. **(A)** Representative liver sections show Oil Red O staining. **(B)** The stained areas were quantified in five randomly chosen microscopic fields per liver section (200× magnification) with Image-Pro software. Data are presented as the means of percent stained areas ± SD (n = 5 mice/time point/genotype; *, *p* < 0.05; **, *p* < 0.01).

### Hepatocyte-specific *Bmal1* knockout leads to suppression of expression peaks in a subset of cell cycle components and regulators in regenerating livers

We found that in *Bmal1*^+/+^ mice subjected to PH, hepatic *Bmal1* mRNA expression fluctuated approximately 5-fold (Fig. 5A). To evaluate Bmal1 transcriptional activity, we quantified transcript levels of hepatic *Dbp*, a direct target gene of Bmal1(40). As a result, in *Bmal1*^+/+^ regenerating livers, *Dbp* mRNA exhibited oscillatory expression with an approximate 4-fold magnitude, which was inversely correlated with *Bmal1* mRNA levels, as anticipated, whereas in *Bmal1*^hep-/-^regenerating livers, *Dbp* mRNA expression lost such oscillation (Fig. 5A). Bmal1 is a suppressor of Per1, Per2, Cry1, and Cry2, which are additional components of the circadian clock (13). The transcript levels of *Per1, Per2, Cry1*, and *Cry2* fluctuated as liver regrowth advanced in *Bmal1*^+/+^ mice. Bmal1 deficiency increased mRNA expression of *Per2* and *Cry1*, but mildly affected *Per1* and *Cry2*, in regenerating livers (Fig. 5A). These data indicated that the circadian clock was fully operating during hepatic regeneration.

**Figure 5.**
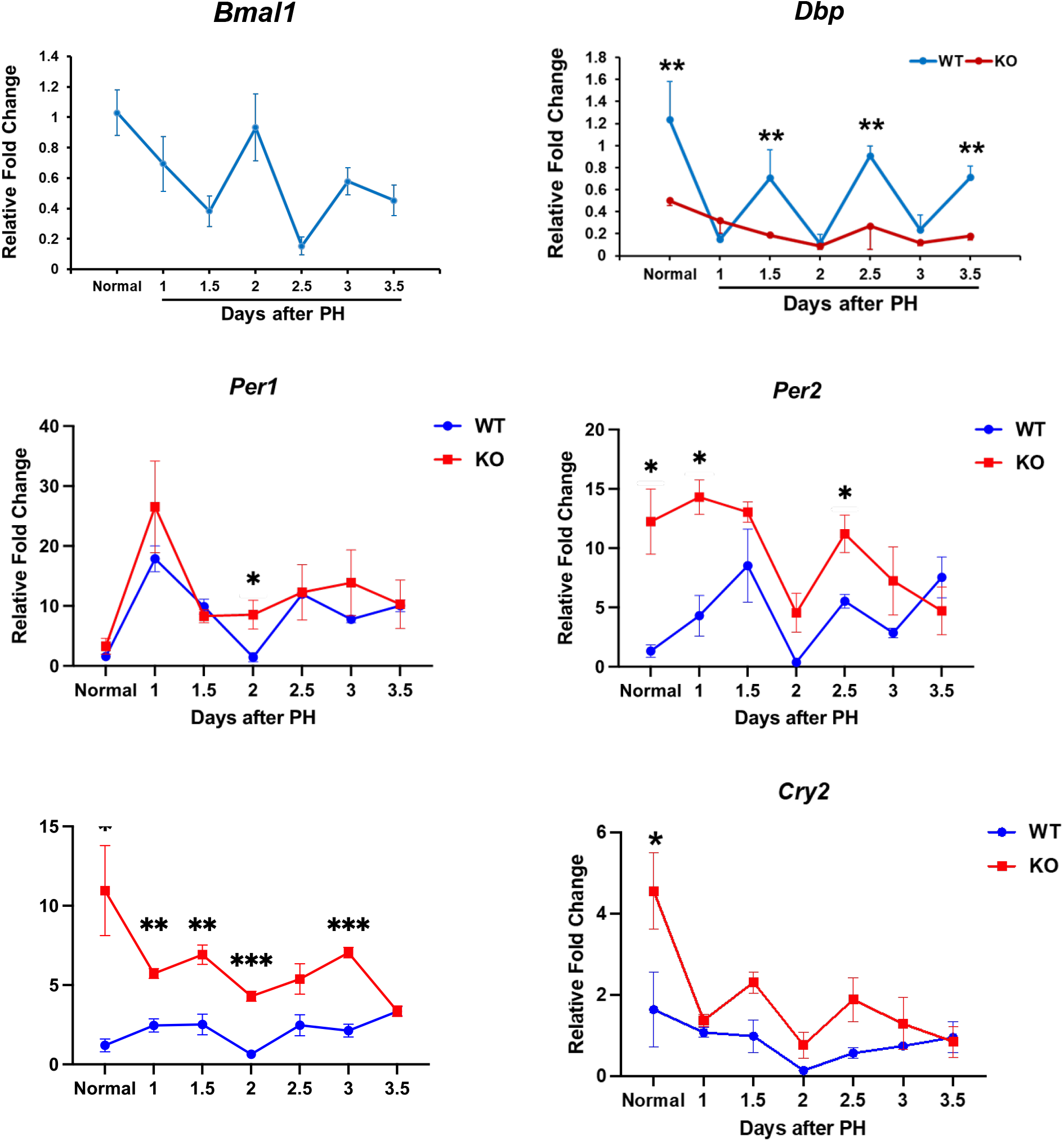
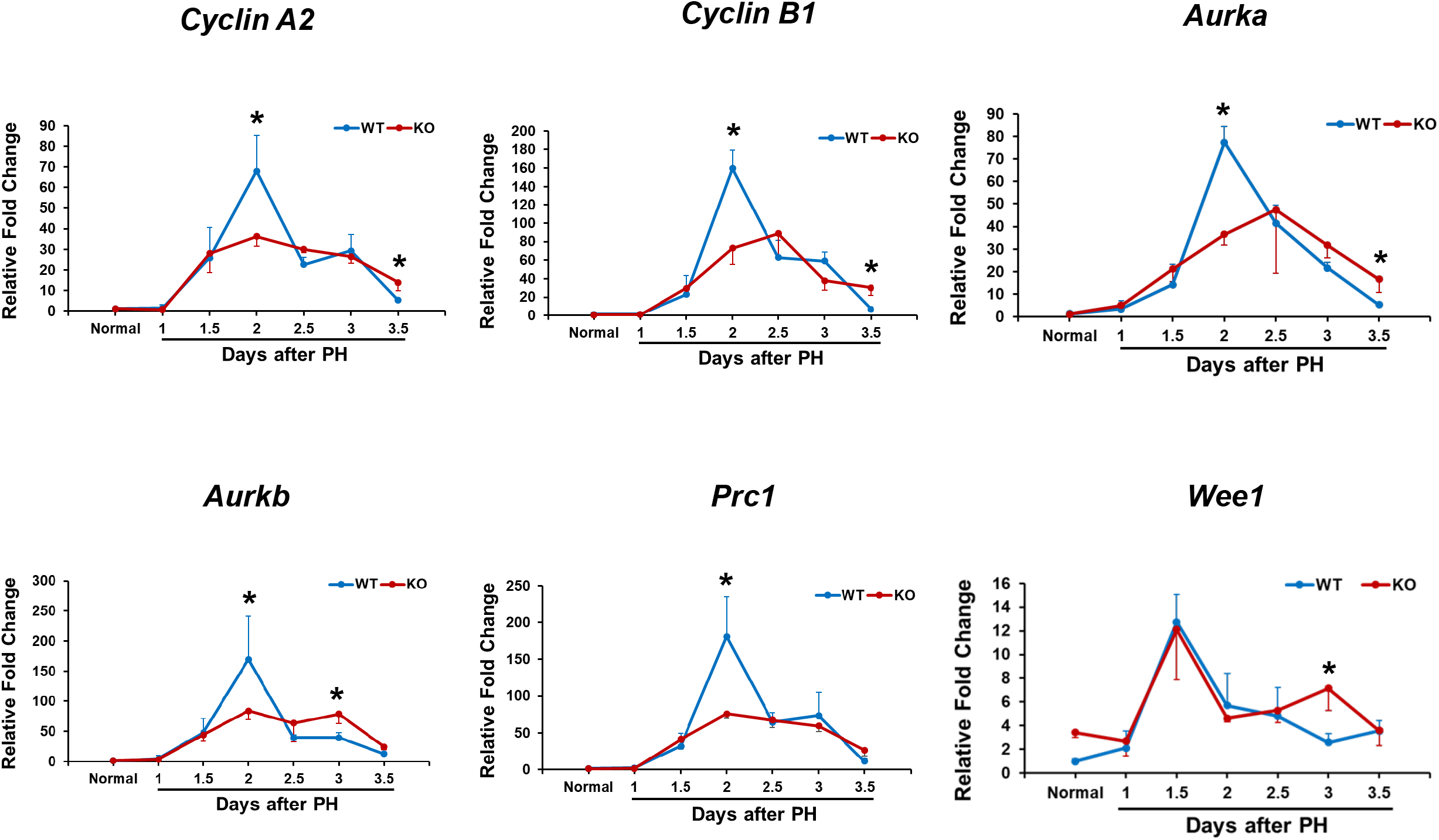
mRNA expression of a subset of circadian clock components and cell cycle components and regulators in regenerating liver. *Bmal1*^+/+^ (WT) and hepatocyte-specific *Bmal1* knockout (KO) 3-4-month-old male mice were subjected to PH and sacrificed at the indicated time points. Total liver RNA samples were prepared and used for quantifying mRNA levels of the genes indicated by RT-qPCR. **(A)** mRNA expression of a subset of circadian clock components. **(B)** mRNA expression of a subset of cell cycle components and regulators. Data are expressed as the mean fold change relative to mRNA levels pre-PH in WT mice ± SD (n = 5 mice/time point/genotype; *, *p* < 0.05; **, *p* < 0.01).).

To examine cell cycle progression in cycling hepatocytes at the molecular level, we assessed the expression of a subset of genes encoding cell cycle components and regulators in regenerating livers (Fig. 5B). In partially hepatectomized *Bmal1*^+/+^ mice, hepatic mRNA expression of *cyclin A2, cyclin B1, aurora kinase A* (*Aurka*), *aurora kinase B* (*Aurkb*), and *protein regulator of cytokinesis 1* (*Prc1*) consistently exhibited peak expression at day 2 post-PH. In the absence of Bmal1, the formation of these peaks was completely abrogated. Thus, Bmal1 is required for these genes to reach peaks in terms of their mRNA expression in regrowing livers. Exceptionally, the formation of the expression peak of the *Wee1* gene, which encodes a kinase controlling cell mitosis, was not dependent on Bmal1 (Fig. 5B). At the protein level, cyclin D1 was persistently expressed from day 1 to day 7 after PH in both liver genotypes. Overall, *Bmal1*^hep-/-^livers expressed more cyclin D1 at most time points post-PH than *Bmal1*^+/+^ control livers (Fig. 6). Notably, the expression of *cyclin A2, cyclin B1*, and *p-Cdc2* was oscillatory and most abundant on days 2 and 4 following PH in *Bmal1*^+/+^ regenerating livers, whereas these oscillations were largely repressed in *Bmal1*^hep-/-^regenerating livers (Fig. 6). Taken together, these data demonstrate that Bmal1 exhibits rhythmic activity and determines the formation of expression peaks of a group of cell cycle components and regulators as liver regrowth advances.

**Figure 6.**
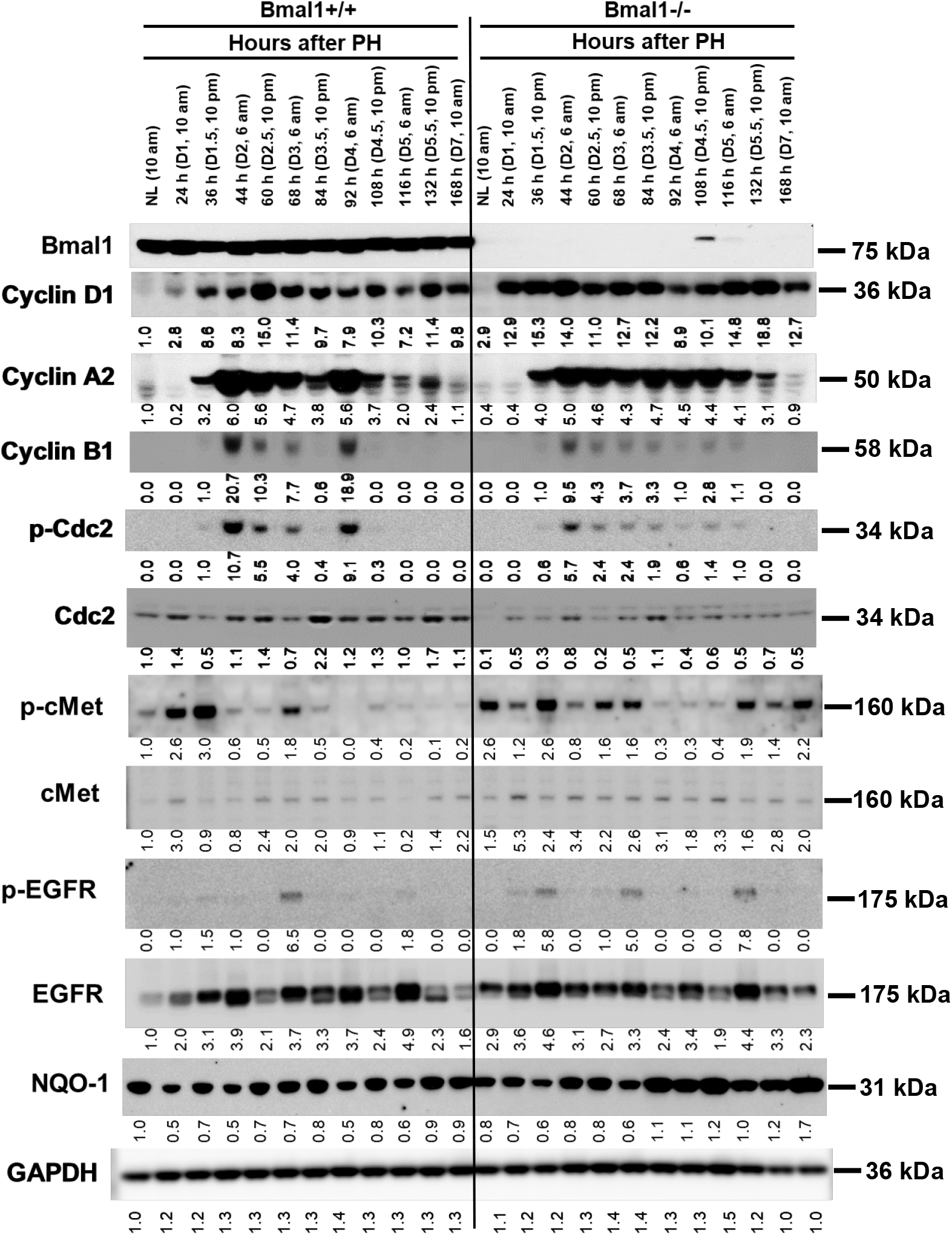
Protein expression of a subset of cell cycle components and mitogenic signaling molecules in regenerating liver. *Bmal1*^+/+^ (WT) and hepatocyte-specific *Bmal1* knockout (KO) 3-4-month-old male mice were subjected to partial hepatectomy (PH) and sacrificed at the indicated time points. Liver lysates prepared from 5 mice per time point per genotype were pooled with equal amount of proteins from each preparation. Western blotting was performed using antibodies against the proteins indicated. Glyceraldehyde 3-phosphate dehydrogenase (GADPH) was used as a loading control. Relative densitometry was normalized against GAPDH. NL, normal liver.

### Hepatocyte-specific *Bmal1* knockout causes dysregulated activation of mitogenic signaling molecules in regenerating livers

cMet and epidermal growth factor receptor (EGFR) are the two most critical mitogenic signaling molecules involved in liver regeneration (24). We found that, in *Bmal1*^+/+^ mice, relative to the pre-PH state, hepatic c-Met displayed increased activity (phosphorylation) during the first 3 days following PH. In *Bmal1*^hep-/-^mice, compared with *Bmal1*^+/+^ mice, hepatic c-Met exhibited markedly (2.6-fold) higher basal activity pre-PH and an altered activation pattern post-PH (Fig. 6). In *Bmal1*^+/+^ regrowing livers, EGFR was activated most prominently at day 3, whereas in *Bmal1*^hep-/-^regrowing livers, this molecule was activated at multiple time points (days 1.5, 3, and 5) and post-PH (Fig. 6). Together, we demonstrated that Bmal1 determines the basal activity of c-Met in homeostatic livers and the timing of c-Met and EGFR activation in regenerating livers.

### Hepatocyte-specific *Bmal1* knockout disrupts the expression patterns of genes involved in lipid metabolism in regenerating livers

We examined the expression patterns of a group of genes known to participate in lipid metabolism in regrowing livers (Fig. 7). Rev-erbα is a state-dependent regulator of liver energy metabolism and serves to buffer against metabolic challenge (15). *Bmal1*^+/+^ mice showed persistently suppressed, whereas *Bmal1*^hep-/-^mice displayed further suppressed, mRNA expression of hepatic *Rev-erbα* following PH. Peroxisome proliferator activated receptor gamma (Pparg) promotes fatty acid uptake and triglyceride synthesis and storage (23). Cd36 is a free fatty acid transporter responsible for the uptake of fatty acid (5). Fatty acid desaturase 1 (Fads1) is involved in long-chain polyunsaturated fatty acid metabolism (18). During the first 1.5 days after PH, hepatic expression of the genes encoding these 3 proteins was inhibited due to the absence of Bmal1. Pparα increases fatty acid uptake, esterification, and trafficking as well as lipid oxidation (4, 23). Fatty acid synthase (Fasn) catalyzes fatty acid synthesis (17). Three days after PH, without *Bmal1*, the transcript levels of hepatic *Pparα, Pparg*, and *Fasn* were all down-regulated. Sterol regulatory element-binding transcription factor 1 (SREBF1) induces <u>lipogenesis</u> in the liver (32). The expression of *Srebf1*, as well as *Fads2*, did not show a Bmal1-dependent difference in regenerating livers. *Elovl6* encodes elongation of very long chain fatty acids protein 6 and is a target gene of Srebf1. The lack of Bmal1 caused a transient upregulation of hepatic Elovl6 mRNA expression on day 2 following PH. Taken together, these results demonstrated that Bmal1 is required for maintaining PH-dependent expression patterns of a subset of genes critical for lipid metabolism.

**Figure 7.**
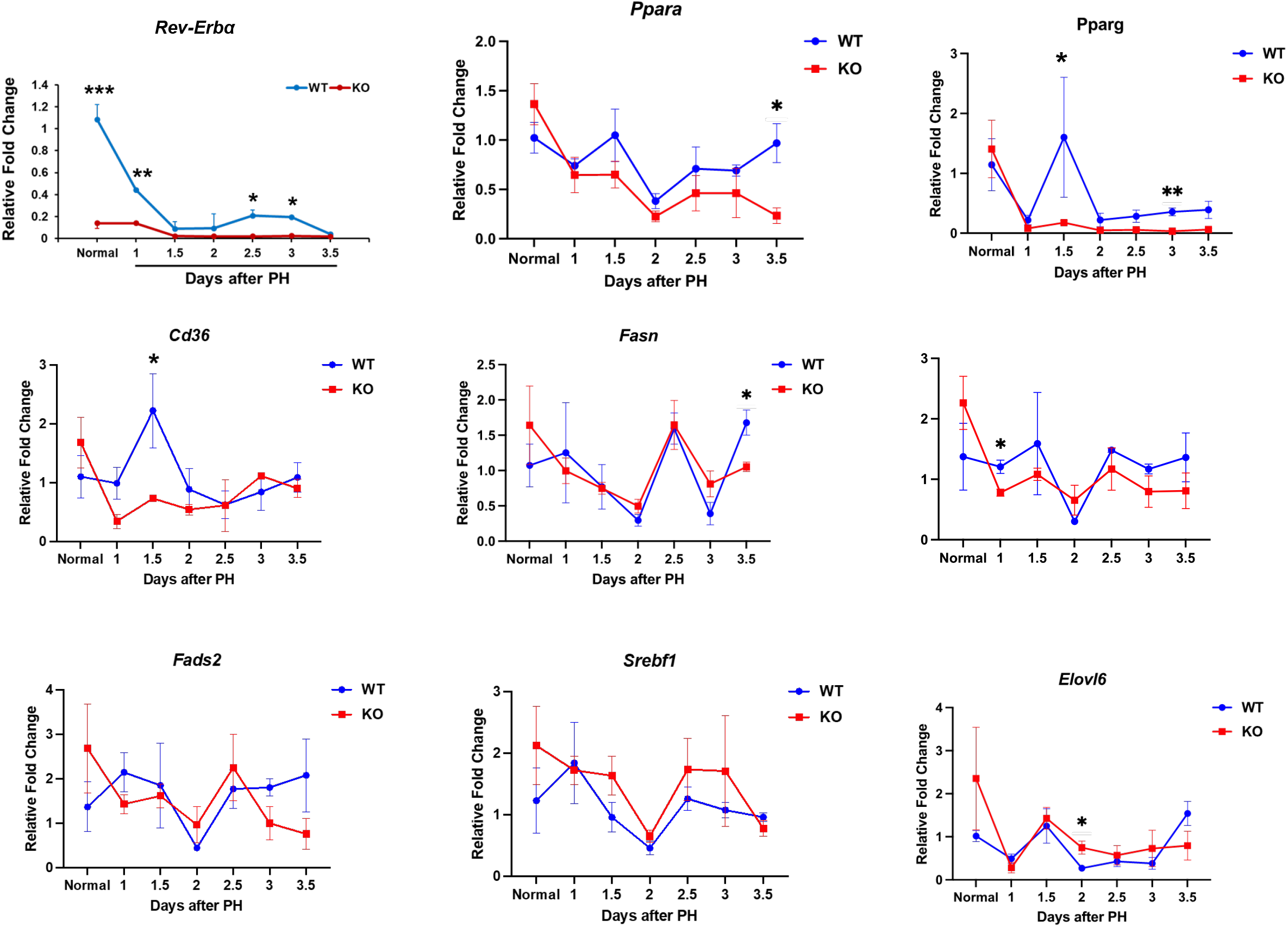
mRNA expression of a group of genes involved in lipid metabolism in regenerating liver. *Bmal1*^+/+^ (WT) and hepatocyte-specific *Bmal1* knockout (KO) 3-4-month-old male mice were subjected to PH and sacrificed at the indicated time points. Total liver RNA samples were prepared and used for quantifying mRNA levels of the genes indicated by RT-qPCR. Data are expressed as the mean fold change relative to mRNA levels pre-PH in WT mice ± SD (n = 5 mice/time point/genotype; *, *p* < 0.05; ***, *p* < 0.001).).

### Hepatocyte-specific *Bmal1* knockout causes Nrf2 activation during late stages of liver regrowth

We calculated liver-to-body weight ratios to estimate gross regrowth of resected livers. The liver-to-body weight ratios were not significantly different between *Bmal1*^+/+^ and *Bmal1*^hep-/-^mice pre-PH (Fig. 8A), consistent with other’s report (26). *Bmal1*^hep-/-^mice regained more liver mass than *Bmal1*^+/+^ mice by day 5.5 post-PH. Overall, the two genotype groups of mice displayed comparable liver mass recovery within the 7 day period post-PH (Fig. 8B). Notably, at day 3.5 and thereafter following PH, *Bmal1*^hep-/-^livers persistently expressed higher levels of NAD(P)H:quinone oxidoreductase 1 (NQO1) than *Bmal1*^+/+^ livers (Fig. 6). NQO1 is a direct target of the transcription factor nuclear factor erythroid 2-related factor 2 (Nrf2), a redox sensor (1). In response to oxidative stress, Nrf2 translocates into the nucleus and transactivates a battery of cellular defense genes (36). Thus, we examined the intracellular distribution of Nrf2 in hepatocytes at day 7 post-PH. As a result, we observed overt nuclear translocation of Nrf2 in *Bmal1*^hep-/-^hepatocytes compared with *Bmal1*^+/+^ hepatocytes (Fig. 8C). These results suggested that absence of Bmal1 in hepatocytes caused oxidative stress as liver regrowth progressed to the later stages, persisting in finally regenerated livers.

**Figure 8.**
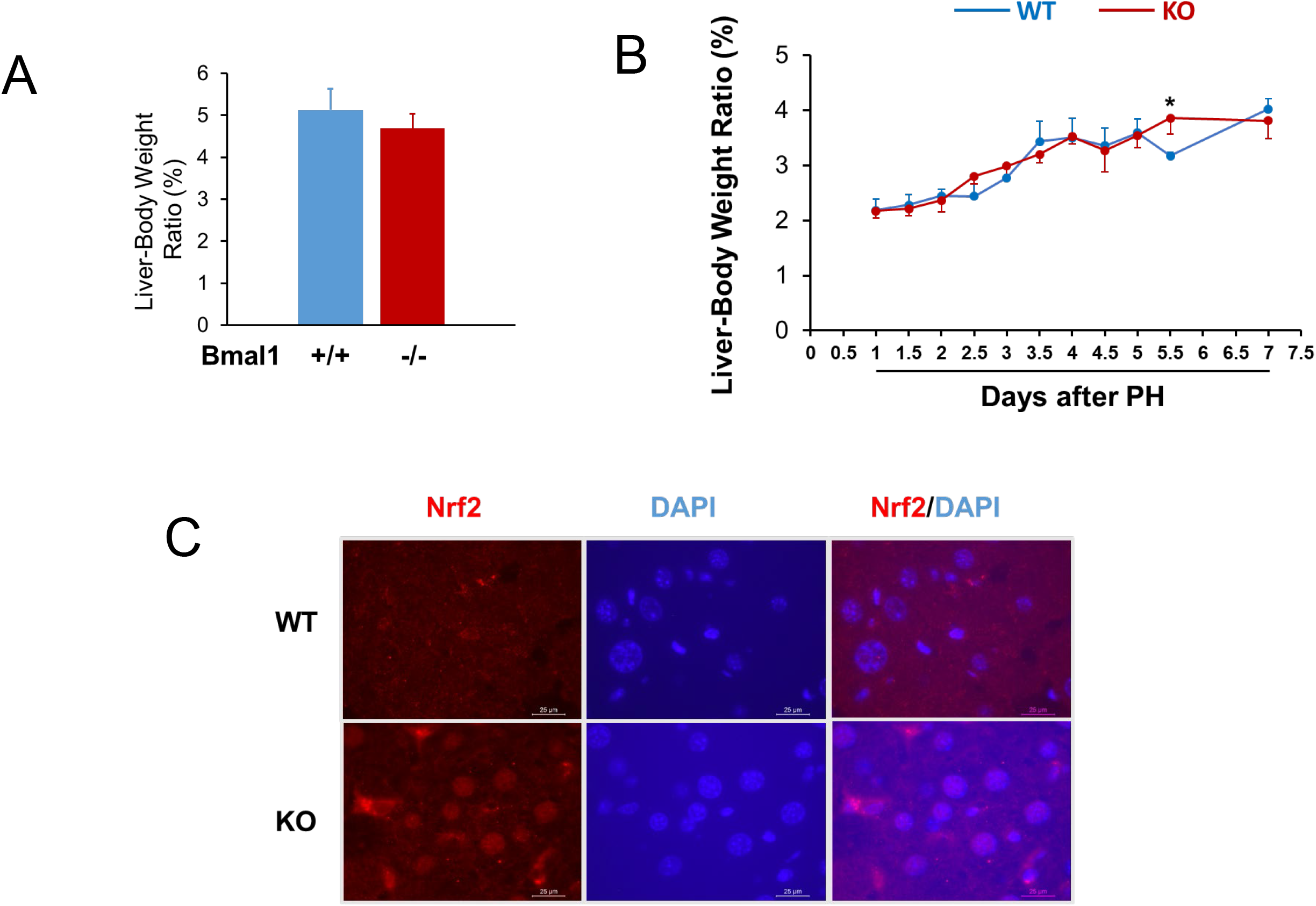
**(A)** Liver-body weight ratios pre-partial hepatectomy (PH). **(B)** Liver regrowth patterns after PH. *Bmal1*^+/+^ (WT) and hepatocyte-specific *Bmal1* knockout (KO) 3-4-month-old male mice were subjected to PH and sacrificed at the indicated time points. Liver and body weights were recorded. Liver-to-body weight ratio was used as a liver regrowth index. The results are presented as mean liver-to-body weight ratio ± SD (n = 5 mice/time point/genotype; *, *p* < 0.05). **(C)** Nrf2 immunofluorescence analysis was performed on liver sections prepared from formalin-fixed and paraffin-embedded liver tissues collected from mice sacrificed at day 7 post-PH. Representative liver sections show Nrf2 (red, cytosol or nucleus) and DAPI (blue, nucleus).

## Discussion

Our studies demonstrated that Bmal1 governs the formation of waves of hepatocyte replication by ensuring timely activation of mitogenic signaling and peak expression of cell cycle regulators and components during liver regeneration. Bmal1 loss of function in hepatocytes caused early entry to the cell cycle, disrupted synchronicity of S-phase progression, and, most strikingly, fully lost rhythmicity of mitosis in cycling hepatocytes in regenerating livers. As early as 36 h post-PH, a significant number of hepatocytes deficient for Bmal1 had already been in the M phase, whereas most hepatocytes sufficient for Bmal1 were only in the first S phase. This initial early entry of hepatocytes into the cell cycle may be due to increased basal activity of c-Met in Bmal1^hep-/-^livers. Here, for the first time, we linked Bmal1 to c-Met activity in the liver under homeostatic conditions. This finding will prompt us and others to delineate how this occurs because dysregulation of the circadian clock is associated with tumor incidence(2). The absence of Bmal1 impaired S-phase progression but did not completely eliminate the formation of DNA synthesis peaks in cycling hepatocytes. This indicated the existence of other agents(s) that more critically control the rhythmicity of DNA synthesis in this setting. Without Bmal1, replicating hepatocytes were unable to form any waves of mitosis in regenerating livers. Thus, these cells rely fully on Bmal1 for timely division. Bmal1 plays an essential role in regulating cyclin A2 and the Cdc2/cyclin B1 pathway. It is well known that the timely accumulation and degradation of cyclin A is required for M-phase progression(25, 34) and that the activity of the Cdc2/cyclin B complex modulates G2/M transition in dividing cells(10). We observed that the expression of cyclin A2, cyclin B1, and p-Cdc2 similarly exhibited large magnitudes of oscillation in *Bmal1*^+/+^ regenerating livers, whereas these oscillations largely disappeared owing to the absence of Bmal1. It was proposed by others that *Wee1* is a direct target gene of Bmal1, and the Bmal1-Clock/Wee1/Cdc2 pathway determines the timing of hepatocyte division in liver regeneration(20). However, we found that the peak formation of *Wee 1* mRNA expression was not affected by the loss of function of Bmal1 in regenerating hepatocytes, which does not support this theory. It is known that c-Met and EGFR together mediate mitogenic signaling essential for hepatocyte proliferation, because their combined loss completely abolishes liver regeneration(24). We found that Bmal1 is required to orchestrate the timed activation of c-Met and EGFR as liver regrowth advances. Aurka, Aurkb, and Prc1 are critical regulators of cell division(11, 16). We showed that Bmal1 is essential for reaching peaks in its expression. Collectively, we propose that Bmal1 centrally coordinates the timing and levels of the activity of mitogenic signaling molecules and the expression of cell cycle regulators and components, thus driving the synchronicity and rhythmicity of the initiation and progression of the cell cycle of cycling hepatocytes during liver regeneration. Of note, a very recent report shows that, in *Bmal1* knockout mice, the skin and liver still exhibit 24-hour oscillations of the transcriptome and proteome without daily light (28). The report proposed that ETS (erythroblast transformation specific) transcription factor family and the redox system may support the oscillations independent of Bmal1. Thus, it is highly likely that these factors may partially compensate for the loss of Bmal1 during liver regeneration, which needs to be further investigated.

Proliferating hepatocytes accumulate fat (steatosis), which is a naturally occurring event, as the liver is regenerating. Excessive lipids provide cellular components for newly regenerated hepatocytes and act as signaling molecules to promote liver regrowth (8). Our previous work demonstrated that cycling hepatocytes rhythmically accumulate fat, which is closely linked to hepatocyte mitosis after PH (42). Here, we revealed that, without Bmal1 in hepatocytes, initial fat deposits were reduced, and subsequent fat accumulation was lost as liver regrowth progressed. Thus, proliferating hepatocytes largely rely on Bmal1 to accumulate lipids. How the steatosis is formed in regenerating liver remains unclear. Our data suggest that Bmal1 coordinates the expression of genes involved in lipid uptake, synthesis, storage, and oxidation. The consequence of loss of such coordination is the reduction or prevention of lipid accumulation in regenerating liver. Of note, a report showed that liver-specific *Bmal1* knockout mice display an increased steatosis phenotype with suppressed *de novo* lipogenesis and fatty acid oxidation in alcohol liver disease (39). Therefore, our findings revealed a liver regeneration-dependent role of Bmal1 in lipid metabolism.

Liver regeneration requires highly coordinated and complex interactions between different liver cell types (21). For instance, after PH, Kupffer cells produce proinflammatory cytokines such as IL-6 to prime hepatocytes to respond to hepatocyte growth factor, which is elaborated by hepatic stellate cells (HSCs), stimulating hepatocytes to enter the cell cycle first. In turn, replicating hepatocytes release growth factors such as platelet-derived growth factor to drive nonparenchymal cell proliferation, which occurs 24 hours later than that of hepatocytes. It is known that the circadian clock operates in liver nonparenchymal cells and regulates their functions (29, 38, 41). Bmal1 controls glycolysis, cell cycle progression, and fibrotic phenotypes in HSCs (38). Bmal1 modulates inflammatory cytokine production, glycolysis, and polarization of Kupffer cells, but the loss of Bmal1 in these cells does not affect pathogenesis of alcoholic liver disease (31, 39, 41). Bmal1 is also expressed in cholangiocytes and regulates their hyperplasia in cholestasis (14, 29). These pieces of evidence support important roles of Bmal1 in liver nonparenchymal cells in homeostasis and disease. Here we demonstrate that the loss of function of Bmal1 disrupts the cell cycle progression and lipid metabolism in hepatocytes. Whether and how liver nonparenchymal cells are affected in this setting needs to be investigated.

Our studies showed that, although hepatocytes deficient in Bmal1 lose cell cycle rhythm following PH, they are still able to expand, eventually recovering the lost liver mass. However, the regenerated livers are in an oxidative stress state, which is manifested by Nrf2 activation during the later stage of liver regrowth. It has been reported that Bmal1 directly regulates Nrf2 expression and activity in β-cells (19), macrophages (9), lung (27), and kidney (35). The lack of Bmal1 causes diminished Nrf2 activity and oxidative stress in these cells and organs. It has also been shown that, in hepatocytes, Nrf2 activation directly upregulates the expression of the clock repressor gene *Cry2*, thereby repressing Bmal1/Clock-regulated E-box transcription (37). These reports outline the integration of the circadian clock with redox balance in various cell types. NQO1 expression is solely controlled by Nrf2 during liver injury and regeneration, and serves as a reliable index of Nrf2 activity (1, 43). We found that, compared to *Bmal1*^+/+^ mice, hepatic Nrf2 activity was slightly lower pre-PH, largely comparable during the first three days after PH, and persistently higher after 3.5 days post-PH in hepatocyte-specific *Bmal1* knockout mice. These observations suggested that oxidative stress in regenerated livers is not directly and fundamentally caused by the lack of Bmal1 in hepatocytes. Instead, it was a new event that occurred following disruption of the hepatocyte cell cycle. Therefore, it is highly likely that this is a consequence of the arrhythmic cell cycle in *Bmal1*-deficient proliferating hepatocytes.

In summary, we demonstrated that, after PH, fluctuating Bmal1 activity partially drives synchronous and rhythmic S-phase progression, and fully controls synchronous and rhythmic M-phase progression of cycling hepatocytes, which eventually restores the redox balance in regenerated livers. Mechanistically, Bmal1 ensures the formation of the expression peaks of a group of cell cycle components and regulators, coordinates the timely activation of mitogenic signaling molecules, and modulates waves of fat accumulation as liver regeneration progresses.

## Notes

### Competing Interest Statement

The authors have declared no competing interest.

